# Samplot: A Platform for Structural Variant Visual Validation and Automated Filtering

**DOI:** 10.1101/2020.09.23.310110

**Authors:** Jonathan R. Belyeu, Murad Chowdhury, Joseph Brown, Brent S. Pedersen, Michael J. Cormier, Aaron R. Quinlan, Ryan M. Layer

## Abstract

Visual validation is an essential step to minimize false positive predictions resulting from structural variant (SV) detection. We present Samplot, a tool for quickly creating images that display the read depth and sequence alignments necessary to adjudicate purported SVs across multiple samples and sequencing technologies, including short, long, and phased reads. These simple images can be rapidly reviewed to curate large SV call sets. Samplot is easily applicable to many biological problems such as prioritization of potentially causal variants in disease studies, family-based analysis of inherited variation, or *de novo* SV review. Samplot also includes a trained machine learning package that dramatically decreases the number of false positives without human review. Samplot is available via the conda package manager or at https://github.com/ryanlayer/samplot.

**Contact:** Ryan Layer, Ph.D., Assistant Professor, University of Colorado Boulder.

## Background

Structural variants, which include mobile elements, deletions, duplications, inversions, and translocations larger than 50bp, can have serious consequences for human health and development [1–3] and are a primary source of genetic diversity [4,5]. Unfortunately, state-of-the-art SV discovery tools still report large numbers of false positives [6–9]. While filtering and annotation tools can help [10,11], tuning these filters to remove only false positives is still quite difficult. As the human eye excels at pattern recognition, visual inspection of sequence alignments in a variant region can quickly identify erroneous calls, making manual curation a powerful part of the validation process [6,12,13]. For example, a recent study of SVs in 465 Salmon samples [6] found that 91% of SVs reported using Illumina paired-end sequencing data were false positives. However, the false positive rate plummeted to 7% (according to long-read sequence validation) subsequent to visual inspection [12]. This study highlights the essential step of removing false positives from SV calls and the effectiveness of visual review to identify the real variants.

Tools such as the Integrative Genomics Viewer (IGV) [14], bamsnap [15] and svviz [13] enable visual review of SVs, but they can be cumbersome or complicated, slowing down the review process and often limiting the number of SVs that can be considered. IGV is optimized for single-nucleotide variant visualization, making it easy to zoom into particular loci to identify base mismatches in read pileups. While IGV can be configured for SV viewing (i.e., viewing reads as pairs, sorting by insert size), visualizing large variants is difficult. The software often loads slowly for large variants which require plotting large numbers of reads. To address slow loading, IGV defaults to sampling a subset of reads and stops displaying alignment data when viewing broad regions, both of which further complicate SV interpretation. IGV has a batch image generation mode for curating many SV calls, but it lacks the full suite of options necessary for SV image optimization. Bamsnap provides a similar visualization optimized for small regions, although review can be faster as static images are created rather than a dynamic viewer as in IGV.

Svviz provides an innovative view of the sequencing data. Alignments are divided into two plots. One plot shows reads that align to the reference allele and the second shows reads that align to the alternate allele created by the SV. Although the clear separation of evidence by reference and alternate alleles is an improvement, svviz plots can be large, complex, and time-consuming to review. Svviz plots also depend on the purported SV breakpoints. Since reads are realigned to a specific alternate allele, any imprecision in the SV breakpoints will affect the visualization making it impossible to differentiate between an absent SV and a slightly incorrect call.

Samplot provides a set of tools designed specifically for SV curation. Samplot’s plotting function creates images designed for rapid and simple, but comprehensive, visual review of sequencing evidence for the occurrence of an SV. The Samplot VCF functionality generates plots for large numbers of SVs contained in a VCF file and provides powerful and easy-to-use filters to refine which SVs to plot, enhancing and streamlining the review process. Finally, the Samplot-ML tool automates much of the review process with high accuracy, minimizing required human-hours for curation.

## Results

Samplot provides a quick and straightforward platform for rapidly identifying false positives and enhancing the analysis of true positive SV calls. Samplot images are a concise SV visualization that highlights the most relevant evidence in the variable region and hides less informative reads. This view provides easily curated images for rapid SV review. Samplot supports all major sequencing technologies and excels at the comparison between samples and technologies. Users generally require less than 5 seconds to interpret a Samplot image [12], making Samplot an efficient option for reviewing thousands of SVs. The simple images contrast with existing tools such as IGV, bamsnap, and svviz which allow more in-depth, but more complex and time consuming, SV-region plotting (see **Figure 1, Supplemental Figures 1-4**).

**Figure 1.**
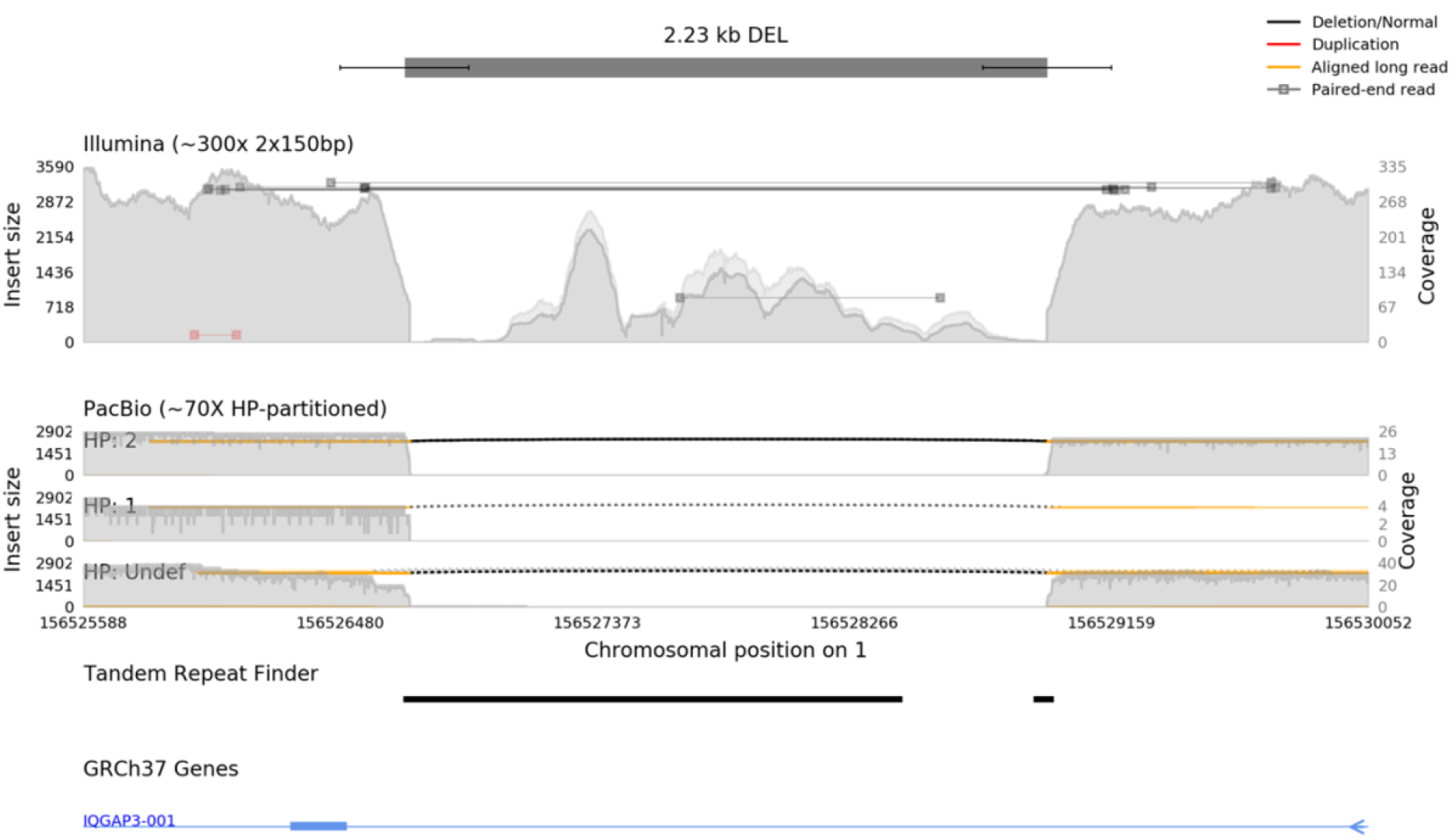
Samplot creates multi-technology images specialized for SV call review. A putative deletion call is shown, with the call and confidence intervals at the top of the image (represented by a dark bar and smaller lines). Two sequence alignment tracks follow, containing Illumina paired-end sequencing and Pacific Biosciences (PacBio) long-read sequencing data, each alignment file plotted as a separate track in the image. PacBio data is further divided by haplotype (HP) into subplots. Reads are indicated by horizontal lines and color-coded for alignment type (concordant/discordant insert size, pair order, split alignment, or long read). The coverage for the region is shown with the grey-filled background, which is split into map quality above or below a user-defined threshold (in dark or light grey respectively). An annotation from the Tandem Repeats Finder^18^ indicates where genomic repeats occur. A gene annotation track shows the position of introns (thin blue line) and exons (thick blue line) near the variant; a small blue arrow on the right denotes the direction of transcription for the gene.

Samplot is also designed for easy application to various types of SV study, such as comparing the same region across different samples (**Supplemental Figure 5**) and sequencing technologies (**Supplemental Figure 6**) for family, case-control, or tumor-normal studies. Annotations such as genes, repetitive regions, or other functional elements can be added to help add context to SV calls (**Figure 1**).

Samplot supports short-read sequencing from Illumina, long-read sequencing from Pacific Biosciences or Oxford Nanopore Technologies, and linked-read sequencing from 10X Genomics. Since genome sequencing technology advances often drive genomic discoveries, Samplot can easily support new sequence types in the future. Samplot works well for most SV types with each of these sequencing technologies and can also plot images without specifying a variant type, enabling review of complex or ambiguous SV types, or non-SV regions.

Producing images that appropriately summarize the evidence supporting an SV without overwhelming the viewer is an intricate task. Samplot includes the three most essential categories of SV evidence: split reads, discordant pairs, and coverage anomalies. To reduce confusion, we distinguish between sequences and alignments. A sequence (also called a read) is a series of nucleotides produced by a short or long-read sequencing platform. An alignment describes how a sequence (or read) maps to the reference genome. Sequences that originate from a region of a sample’s genome that does not include an SV will have a single complete continuous alignment. When a sequence includes an SV, it will produce multiple alignments or unaligned segments. The configuration of these alignments indicates the SV type. Deletions create gaps between alignments, duplications create overlapping alignments; inversions produce alignments that switch between strands, etc.

An SV in the unsequenced region between the paired-end sequencing reads will have a discordant alignment whose configuration similarly indicates the SV type. Samplot identifies, color-codes, and elevates split or discordant alignments so that users can clearly and quickly distinguish between normal reads and reads supporting different SV types (**Figure 2**). These plots often include scatterings of misaligned reads that can fool automated tools. A visual review can generally quickly determine whether or not groups of reads support an SV, allowing rapid high-confidence variant review.

**Figure 2.**
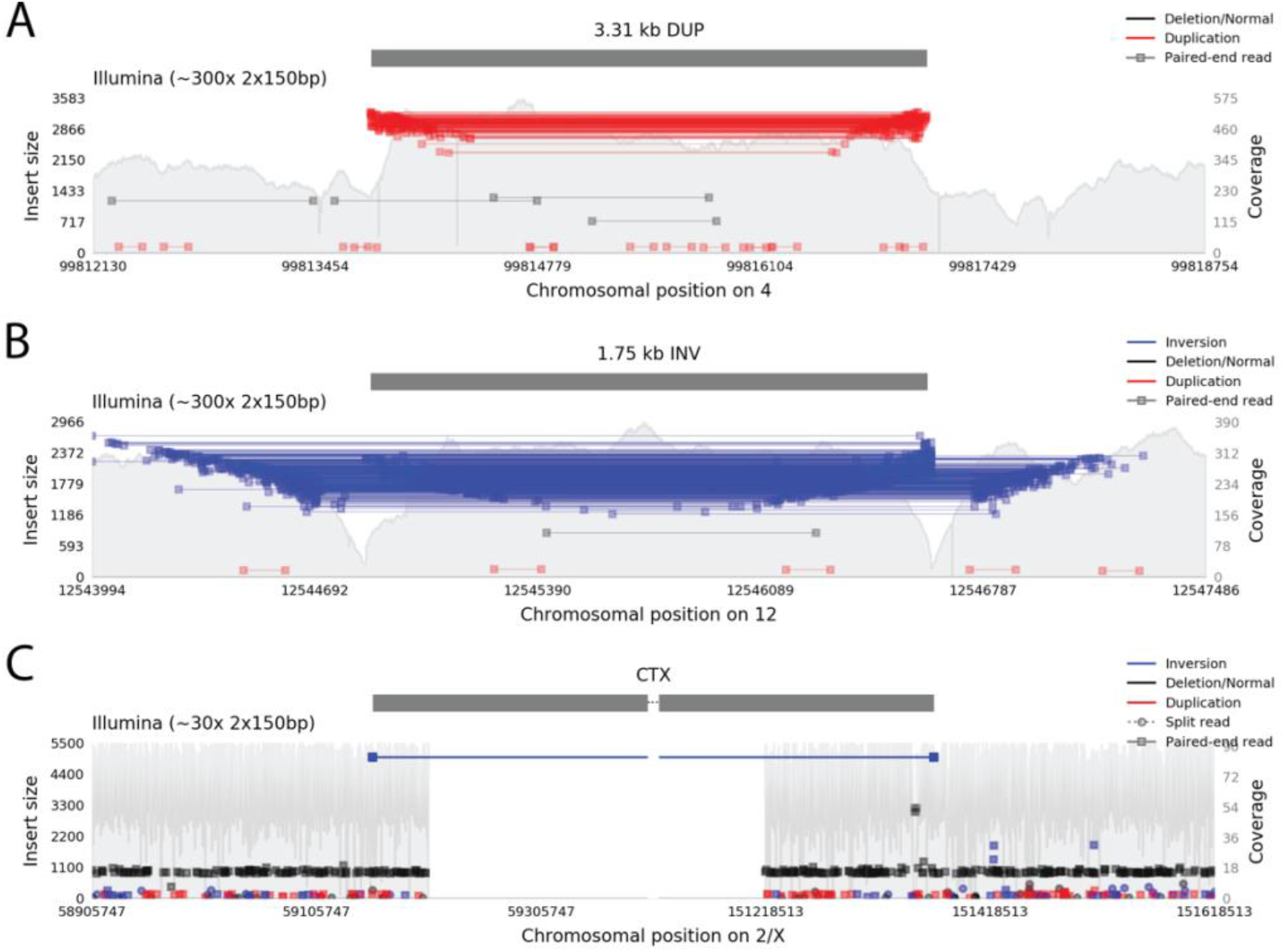
Samplot images of duplication, inversion, and translocation variants. **A)** A duplication variant plotted by Samplot with Illumina short-read sequencing evidence. Reads plotted in red have large insert sizes and inverted pair order (reverse strand followed by forward strand instead of forward followed by reverse), indicating potential support for a duplication. **B)** An inversion variant, with Illumina sequencing evidence. Reads plotted in blue have large insert sizes and same-direction pair alignments (both reads on forward strand, or both on reverse strand). **C)** A translocation variant, with Illumina sequencing. Discordant pairs align to each breakpoint. The blue color of the reads and extremely large insert sizes of these grouped discordant pairs indicate a large inverted translocation.

Coverage depth is also an essential piece of data for evaluating the SVs that affect genomic copy number (copy number variants or CNVs) and can, in some cases, provide the best signal of a CNV. Samplot includes a background track with up to base-pair resolution of the fluctuations in coverage depth across the plot region. Samplot follows a minimal decision-making strategy and makes no computational attempt to assign reads or coverage deviations to putative variant coordinates; this task is left instead to the user via visual curation.

Samplot is implemented in the Python language and utilizes the pysam [16] module to extract read information from alignment (BAM or CRAM) files, then plots reads for review in static images. Speed is a key goal of Samplot, in keeping with the overall focus on simple and rapid SV review, so plots are created using the Matplotlib library, which has been optimized for rapid creation of high-quality images.

### Filtering and viewing SV call sets with Samplot VCF

When working with large SV call sets, especially multi-sample VCF files, users often need to review evidence for SVs in multiple samples together. Samplot provides a VCF-specific option to interrogate such call sets using cohort genotypes, an optional pedigree file in family-based cohorts, and additional annotation fields for filtering and plotting multiple SVs across multiple samples. This enables users to focus on rare variation, variants in certain genome regions, or other criteria related to a research goal. A simple query language that is inspired by slivar [17] allows users to customize filters based on variant annotations in the VCF file. From the chosen variants, a web page is dynamically created with a table of variant information, additional filtering options, and quick access to Samplot images for visual review (**Figure 3**).

Samplot VCF can be readily adapted to experimental needs common in SV studies. For example, a team attempting to identify a causal SV in a familial rare disease study might include a small number of control samples as well as the affected family and use built-in filtering options to plot only variants which appear uniquely in the offspring, with controls included in the resulting images for comparison. Samplot VCF is equally well-suited for other problems such as cohort-based analysis of common SVs or tumor-normal comparison (potentially with multiple samples in each category).

**Figure 3.**
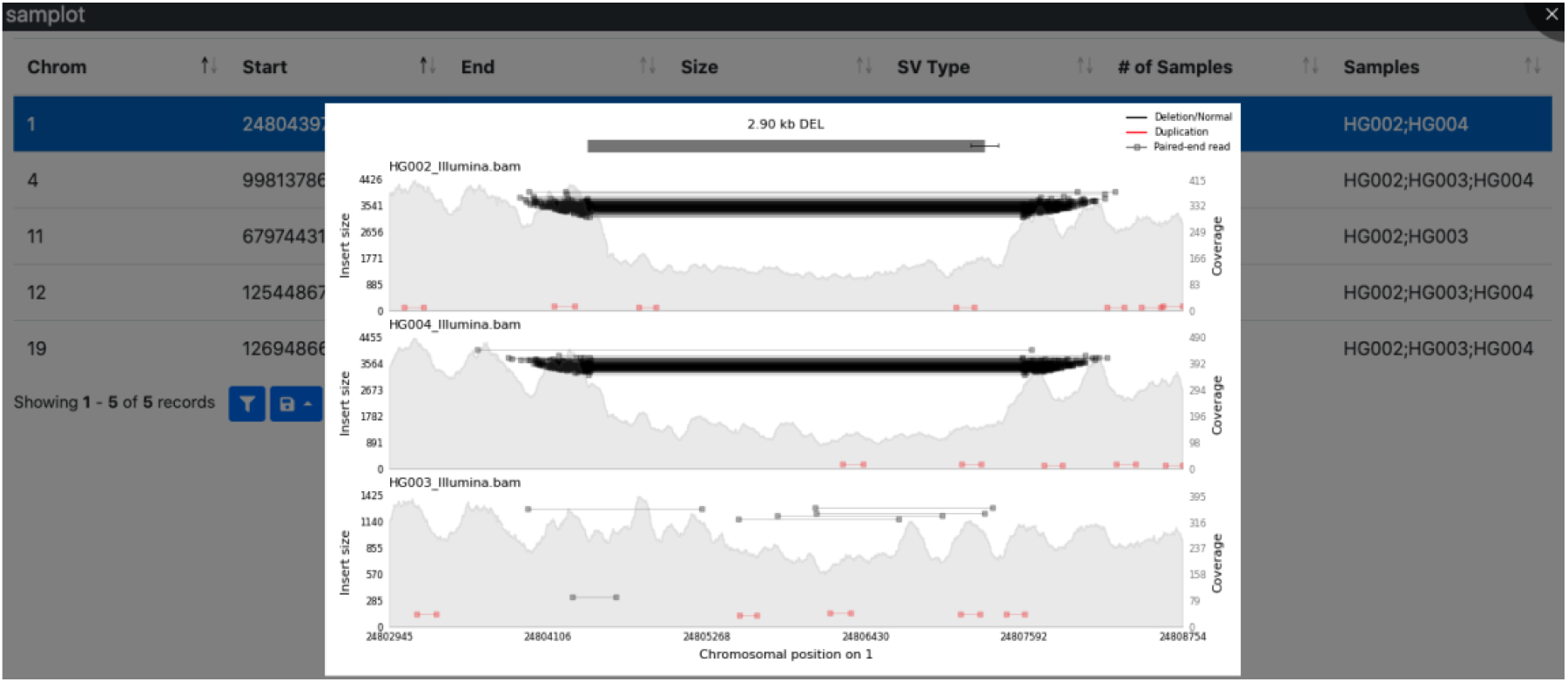
Samplot creates images for quick review of SV VCF files. Samplot’s ‘samplot vc? command will plot all SVs in a VCF file or filter to a subset via user-defined statements. ‘Samplot vcf’ creates an index page and sends commands to ‘samplot plot’, which generates images for each variant that passes the filters. The index.html page displays a table of variant info. Clicking on a row loads a Samplot image, allowing additional filtering or variant prioritization.

### Automated SV curation with Samplot-ML

Convolutional neural networks (CNNs) are an effective tool for image classification tasks. Since Samplot generates images that allow the human eye to adjudicate SVs, it motivated us to test whether a CNN could discern the same patterns. To that end, we developed Samplot-ML, a CNN built on top of Samplot to classify putative SVs automatically. Samplot-ML currently supports deletions. As additional data becomes available, Samplot-ML will support other SV classes as well. The workflow for Samplot-ML is simple: given a whole-genome sequenced sample (BAM or CRAM [18]) as well as a set of putative deletions (VCF [19]), Samplot-ML re-genotypes each putative deletion using the Samplot-generated image. The result is a call set where most false positives are flagged.

Using Samplot-ML, we demonstrate a 51.4% reduction in false positives while keeping 96.8% of true positives on average across short-read samples from the Human Genome Structural Variation Consortium [20]. We also trained a proof-of-concept long-read model with the same architecture and reduced false positives by 27.8%. We expect the long-read performance to improve as more data becomes available. Our model is highly general and can classify SVs in sequences generated by libraries that differ in depth, read length, and insert size from the training set. The Samplot-ML classification process is completely automated and runs at about 100 SVs per second using a GPU and 10 SVs per second using only a CPU. Most SV call sets from methods such as LUMPY [21] and MANTA [22] running on a single genome that yield between 7,000 and 10,000 SVs will finish in about one minute. The result is an annotated VCF with the classification probabilities encoded in the FORMAT field.

While Samplot-ML inherently supports any SV type, the current model only includes deletions. There are too few called duplications, insertions, inversions, and translocations in the available data to train a high-quality model. For example, the 1,000 Genomes Project phase 3 SV call set [4] included 40,922 deletions, 6,006 duplications, 162 insertions, 786 inversions, and no translocations. We expect this limitation to be temporary.

To evaluate the short-read model, we considered the samples from the HGSVC with long-read-validated SVs. First we called SVs in HG00514, HG00733, and NA19240 using LUMPY/SVTYPER [10] (via smoove [23]) and MANTA. Next we filtered those SVs using the heuristic-based method duphold [11], a graph-based SV genotyper Paragraph [24], a support vector machine classifier SV2 [25], and our CNN. In each case, we measured the number of true positives and the number of false positives with respect to the long-read validated deletions using Truvari [26] (**Figure 4A-C**, **Supplemental Table 1**). In all cases both duphold and Samplot-ML removed hundreds of false positives while retaining nearly every true positive. Paragraph and SV2 remove most of the false positives but retain far fewer true positives. Paragraph, similar to other graph-based methods, is also highly sensitive to breakpoint precision (**Supplemental Figure 7**), which explains the differences in its performance between LUMPY and MANTA calls. On average, duphold reduces the number of false positives by 32.2% and reduces true positives by 1.1%. Samplot-ML reduces false positives by 53.4% and true positives by 2.4%. Paragraph and SV2 reduce false positives and true positives 62.5% and 29.9%, and by 84.2% and 63.4%, respectively. A more refined analysis that evaluates the performance by genotype could measure the extent to which the model learns one copy and two copy loss states, but this truth set did not include genotypes. The Genome in a Bottle (GIAB) truth set [27] discussed next had genotypes, and one- and two-copy loss results are decomposed below.

**Figure 4.**
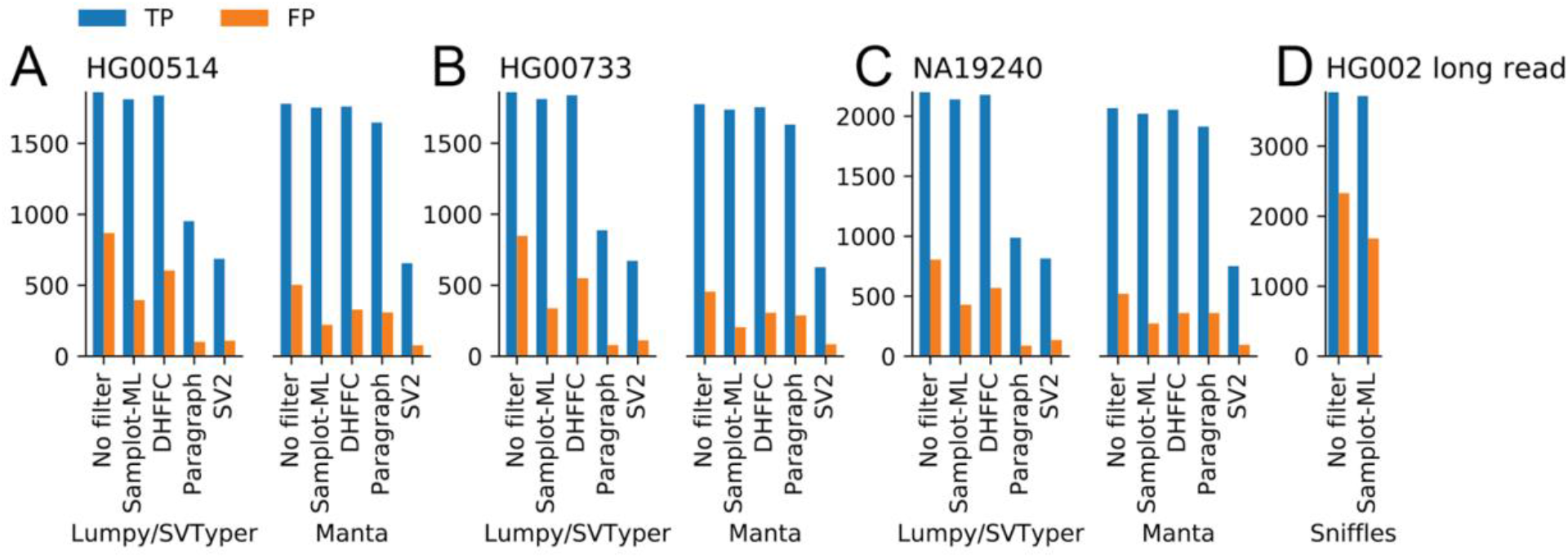
SV filtering performance of duphold (DHFFC) and Samplot-ML. **A-C)** Short-read SVs callsets generated by LUMPY/SVTYPER and MANTA were then filtered by Samplot-ML, duphold (DHFFC), Paragraph, and SV2 were then compared to the long-read-validated truthset. **D)** Long-reads SVs were called with Sniffles, filtered by Samplot-ML, then compared to the GIAB truth set.

The long-read model uses the same architecture and process as the short-read model, except it is trained on genomes sequenced using PacBio Single Molecule, Real-Time (SMRT) Sequencing. Given the limited number of long-read genomes available for training, we consider the model a proof-of-concept that will improve as more long-read genomes become available Since training used the HGSVC samples, the evaluation is based on the GIAB truth set [27] which includes multiple validations, including visual review, for long-read sample HG002. We called SVs using Sniffles [28], filtered those SVs using the CNN, and measured the number of true positives and false positives with Truvari (**Figure 4D**, **Supplemental Table 1**). Samplot-ML reduces false positives by 27.8% and true positives by only 1.4%.

Generality can be an issue with machine learning models. A distinct advantage of training and classifying with Samplot is that its images are relatively consistent across different sequencing properties and the models still perform well when using different sequencing libraries. For example, our short-read model was trained on paired-end sequences 20X with 150 base pair (bp) reads and a 400 bp insert size and the samples in the evaluations above (**Figure 4A-C**) had shorter reads (126 bp reads), a large insert size (500bp), and were sequenced at greater depth (68X). Additionally, we considered two libraries from the same Genome in a Bottle sample (HG002), where one was sequenced at 20X coverage with 150bp reads and 550bp insert size and other was sequenced at 60X coverage with 250bp reads and a 400 bp insert size (**Figure 5A**). The model performed equally well across all libraries, clearly demonstrating that new models are not required for each library. Additionally, between LUMPY and Manta, Samplot-ML correctly genotyped 91.28% of hemizygous deletions (1 copy losses) and 97.26% of homozygous deletions (2 copy losses) for the 20X run (**Figure 4A**). For the 60X run (**Figure 4B**), Samplot-ML correctly genotyped 94.57% of hemizygous deletions and 97.26% of the homozygous deletions. These results clearly show that the model has learned both copy-loss states.

**Figure 5.**
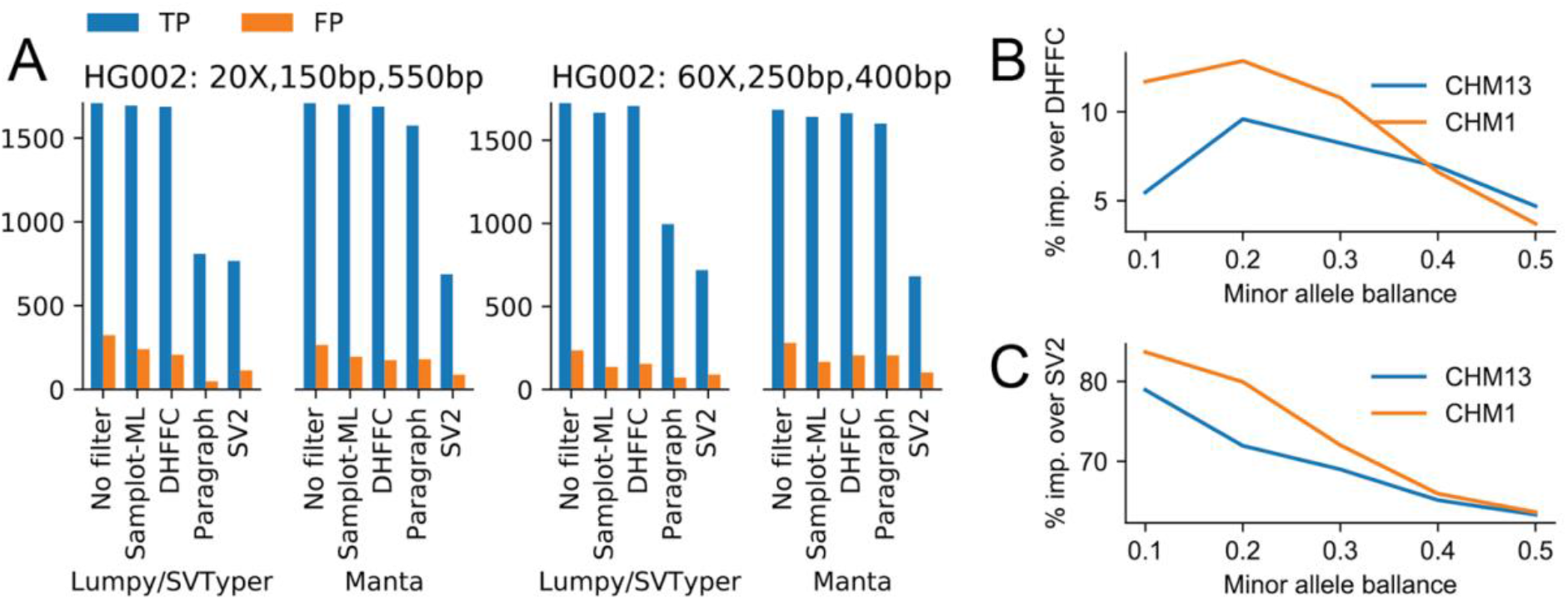
Model performance in data sets that differ from the training set. **A)** The number of true positive and false positive SVs from different SV calling and filtering methods considering the same sample (HG002), sequenced using two libraries with different coverages, read lengths, and insert sizes. **B,C)** The percent increase in true positive SVs that Samplot-ML recovers versus duphold **(B)** and SV2 (**C**) for SVs in simulated mixtures of samples (CHM13 and CHM1 cell lines) at different rates.

Calling SVs in tumor samples can be a challenge when subclones and normal tissue contamination produce variants with a wide range of allele balances (the ratio of reads from the variant allele to the total number of reads). The result is fewer discordant alignments and a less distinct change in coverage, which has a direct effect on the Samplot images (**Supplemental Figure 8**). To test how well our model performs in these instances, we mixed sequences from two diploid cell lines (CHM1 and CHM13) at different rates (**Figure 5B**) then reclassified SVs from a truth set [8] using duphold, SV2, and Samplot-ML. Paragraph was omitted from this experiment due to unresolved runtime errors. For each combination, we compared how many true positive SVs each method recovered from the minor allele. While the recovery rates between the Samplot-ML and duphold were similar, ranging from over 70% when the samples were equally mixed (0.5 allele balance) to less than 40% when the SV minor allele was at 0.1 (**Supplemental Table 2**), Samplot-ML provided an improvement over duphold especially as the minor allele became more rare, peaking at a 12.9% improvement when CHM1 was the minor allele at 20%. SV2’s low sensitivity resulted in poor performance as the minor allele balance decreased. Low-frequency SVs are clearly difficult to detect and filter, but Samplot-ML’s classifier is robust to evidence depth fluctuations, further proof of the generality of the model.

## Discussion

Samplot provides a fast and easy-to-use command-line and web interface to visualize sequence data for most structural variant classes. Pre-screening large SV call sets with Samplot allows researchers and clinicians to remove SVs that are likely to be false-positives and focus orthogonal molecular validation assays on smaller groups of variants with far more true-positives. Rapid review could improve SV detection sensitivity in, for example, low cover sequencing experiments and genomic regions that are thought to be enriched for false positives and excluded in most SV analysis.

We have also trained a convolutional neural network to assist in SV curation. Key to the performance of our model was identifying and training on realistic negative examples (false positives SV calls). In genome feature detection broadly, and SV detection specifically, negatives far outnumber positives. To achieve maximum classification performance, collecting negative training examples must be given as much consideration as any other aspect of the machine learning architecture. Just as it is highly unlikely that any genomic detection algorithm would return a random genomic region as a putative event, we cannot expect that randomly sampled areas of the genome that do not overlap true positives will be good negative examples. Special care must be taken to sample from regions enriched with edge cases that pass detection filters but do not contain true positives. By incorporating putative false positive areas of the genome, we were able to improve the performance of Samplot-ML immensely because these regions strongly resembled the types of false positives that were being made by SV callers.

Our model performs well across sequencing libraries and SV calling algorithms, but currently only supports deletions. As more SV data becomes available, we will extend our model to consider other SV classes. By enabling scalable and straightforward SV review, Samplot can extend robust SV discovery and interpretation to a wide range of applications, from validating individual pathogenic variants to curating SVs from population-scale sequencing experiments.

## Conclusions

Extremely high false-positive SV call rates make rapid curation of callsets a problem of paramount importance for numerous research questions. Samplot and the Samplot-ML classifier together provide powerful, yet simple-to-use tools to curate large SV callsets for high-confidence identification of real SVs. These tools will be widely useful for users seeking to better understand the structure of the human genome and will be especially important as the scientific community sharpens its focus on the impacts of SVs on health, personalized medicine, and diversity.

## Methods

### Samplot-ML model and image generation

Samplot-ML is a resnet [29]-like model that takes Samplot images of putative deletion SVs as input and predicts a genotype (homozygous reference, heterozygous, or homozygous alternate). Samplot-ML was built using Tensorflow [30] and is available at https://github.com/mchowdh200/samplot-ml. For additional model details see **Supplemental Figure 8**. Train and test images were generated using the command:

~~~
samplot plot -c $chrom -s $start -e $end --min_mqual 10 -t DEL -b $bam -o $out file -r $fasta
~~~

Additionally, for SVs with length > 5000 bases, we added `--zoom 1000` which only shows 1000 bp centered around each breakpoint. After an image is generated, we crop out the plot text and axes using imagemagik [31]. Finally, before input into a Samplot-ML model, the vertical and horizontal dimensions are reduced by a factor of eight. Instructions for how to run Samplot-ML can be found in the Supplement under Running Samplot-ML.

### Training Data

#### Short read model

The short read version of Samplot-ML was trained on data from the 1000 Genomes Project (1kg) [4], including the phase three SV call set and the newer high coverage alignments (see Supplemental Tables 3-4 for data URLs). We excluded individuals present in or directly related to individuals in our test sets (NA12878, NA12891, NA12892, HG00512, HG00513, HG00731, HG00732, NA19238, NA19239). While direct relatives of our test set were removed from our training set, it is still possible that samples in our test set contributed some information to the samples in our training in the joint variant calling and genotyping process.

#### True positive Regions

Heterozygous and homozygous deletions were sampled from the GRCh38 liftover of the phase 3 integrated SV map. Although this set contains high confidence SV calls, there were still a few regions that did not exhibit drops in coverage (i.e., false positive call). To minimize the possibility of sampling a false positive, we filtered this set using Duphold’s DHFFC metric which measures the fold change in coverage between the called and flanking regions. To filter, we removed regions with a DHFFC > 0.7. After filtering, we sampled 150,000 heterozygous deletions and 50,000 homozygous deletions.

#### True negative regions

Care must be taken to sample “true negatives” properly. Before choosing a negative set, we must consider the use case of our model. In practice our model will remove false positives from the output set of an SV caller or genotyper. That means that our model will encounter two different classes of regions: those containing real SVs and edge cases that confuse the SV caller’s filters. While we could have sampled regions from homozygous reference samples in the 1kg calls (i.e., samples without deletions) to get “true negatives”, these regions would have had very few discordant alignments and level depths of coverage. Crucially, they would look nothing like the regions that we would want our model to filter.

We took a more principled approach to pick true negatives. Many SV callers have the option to provide a set of “exclude regions”, which prevents the caller from considering potential problematic regions of the genome [32]. Since these are enriched for false positives, we used these regions’ calls as our true negatives. To get variants in these regions, we recalled SVs on the 1kg high coverage alignments using LUMPY [21] with SVTyper [10]. We then selected areas in the resultant calls that intersected problematic regions. To ensure that no true positives were selected, we filtered out regions with a DHFFC ≤ 0.7. Finally, to construct our set of true negatives, we took roughly 35,000 “exclude regions” and 15,000 homozygous reference regions from the 1kg SV call set. One issue with this strategy is that while these regions are enriched for false positives, they could also be enriched for a specific subclass of real variation. We are exploring other training strategies, including using regions that appear to be mutated from short-read data but are normal in long-read data.

#### Long read model

For the long-read model training data, we used PacBio samples from the HGSV consortium (cite) that were present in the 1kg phase 3 SV callset (HG00513, HG00731, HG00732, NA19238, NA19239, see **Supplemental Tables 3-4** for data URLs). This reduced set of samples yielded 5404 true positive regions. Just as with the short-read model, we sampled a mix of “exclude regions” and normal homozygous reference regions. Using the same set of regions called by LUMPY and SVTyper in the short-read alignments, we sampled 452 exclude regions and 4354 homozygous reference regions.

### Training Procedure

From our training set, we held out regions from chromosome 1, 2, and 3 to use as a validation set during training. To train our model, we used stochastic gradient descent with warm restarts (SGDR [33]). The initial learning rate was 0.2 and decayed with a cosine annealing schedule. The initial restart period was set to two epochs and doubled after each restart. We trained for 50 epochs and kept the model with the best validation loss after training was completed.

### Model Testing

#### Short read model

To evaluate the efficacy of the short-read model, we called deletions using both LUMPY/SVTyper and manta on each of our test samples. We then filtered both LUMPY and Manta callsets with Duphold (rejecting calls with DHFFC ≤ 0.7) and Samplot-ML. To compare the filtered callsets with their respective gold standard VCFs (**Supplemental Table 4)**, we used Truvari [26], which compares regions in VCFs based on percent overlap as well as breakpoint accuracy. We used the following truvari command:

~~~
truvari -b $truth_set -c $filtered call set -o $out_dir --sizemax 1000000 --sizemin 300 --sizefilt 270 --pctovl 0.6 --refdist 20
~~~

#### Long read model

To evaluate the long read model, we called deletions using Sniffles [28] on the PacBio HG002 alignments (**Supplemental Table 3**) and filtered the result using Samplot-ML.

#### Variable allele balance simulation

We used sequencing data from human Hydatidiform mole samples CHM1 and CHM13 (See **supplemental Table 4**). Alignments (bams) were generated with BWA-MEM [34] and duplicates were removed with Samblaster[35]. We then randomly subsampled both alignments at a rate of 10 to 90 percent with 10 percent increments and merged CHM1 and CHM13 subsampled alignments such that each mixture added up to 100 percent. We sampled regions (**Supplemental Table 4**) for evaluation that contained homozygous deletions in one sample but not the other. Regions below 10x coverage after filtering reads with less than 10 mapping quality in the non-variant sample were omitted.

#### Paragraph shifted breakpoint experiment

To evaluate paragraphs sensitivity to SV breakpoint precision, we used the HG002 60x 2×150 manta callset VCF and generated VCFs with ±5 base pair shift in start position of each region and ran paragraph for each resulting VCF and evaluated the F1 score using Truvari in the same manner as with the original Samplot-ML model evaluations.

