## Supplemental Figures for "Samplot: A Platform for Structural Variant Visual Validation and Automated Filtering"

### Samplot: multi-technology structural variant sequence data visualization Supplemental Figures

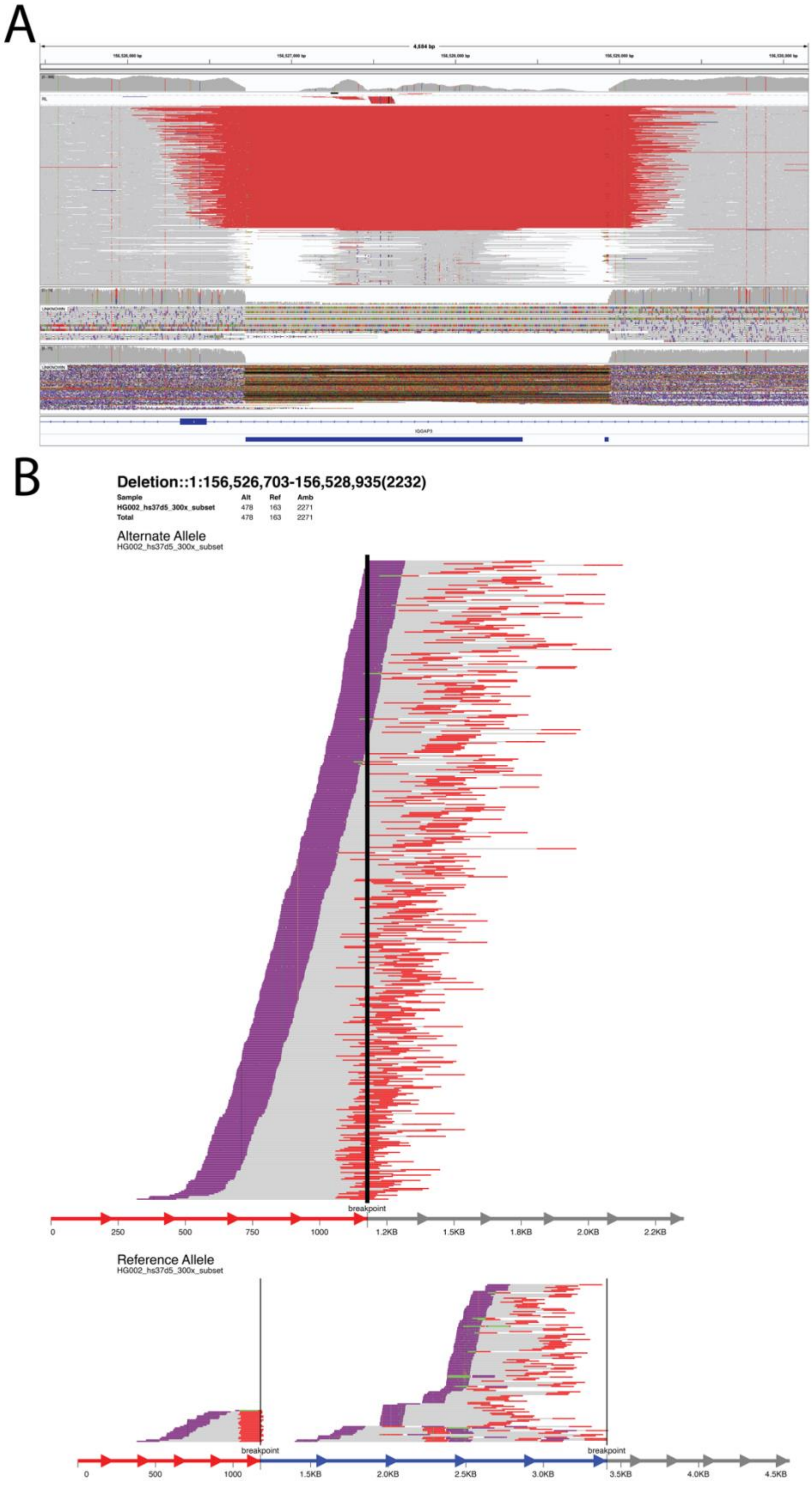


**Supplemental Figure 1. Integrative Genomics Viewer and svviz deletion plots**. **A)** An IGV screenshot of the same deletion variant as shown in Figure 1. Reads are shown as pairs and sorted by insert size, with coverage shown at top of image. **B)** Svviz plot of the same deletion. Reads supporting the alternate allele are shown at top, with reads supporting the reference allele at the bottom. Breakpoints are indicated by dark vertical bars.

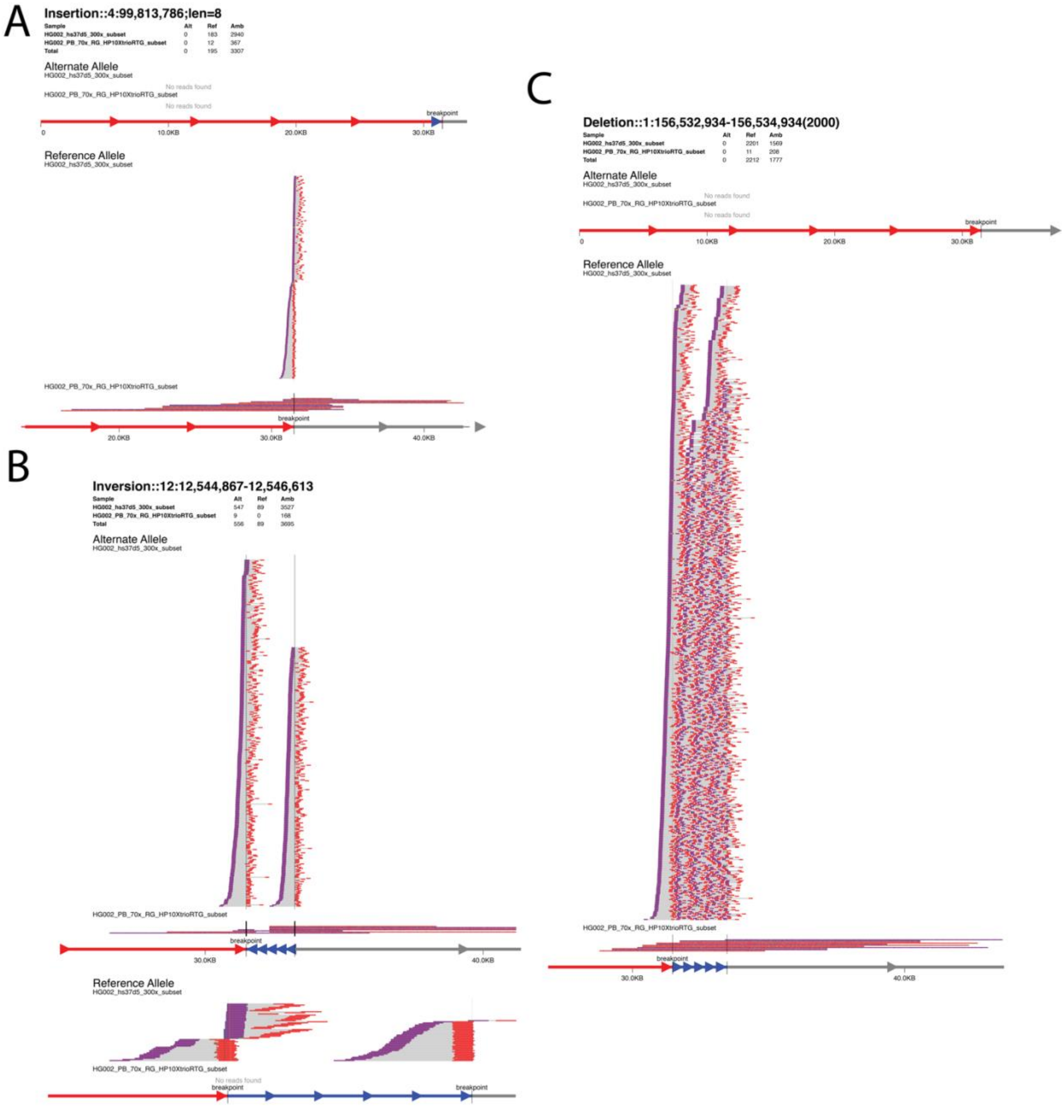


**Supplemental Figure 2. Svviz images for multiple region types. A)** The duplication SV from Figure 2 plotted with svviz. B) The inversion SV from Figure 2 plotted with svviz. **C)** A region with no SV plotted with svviz.

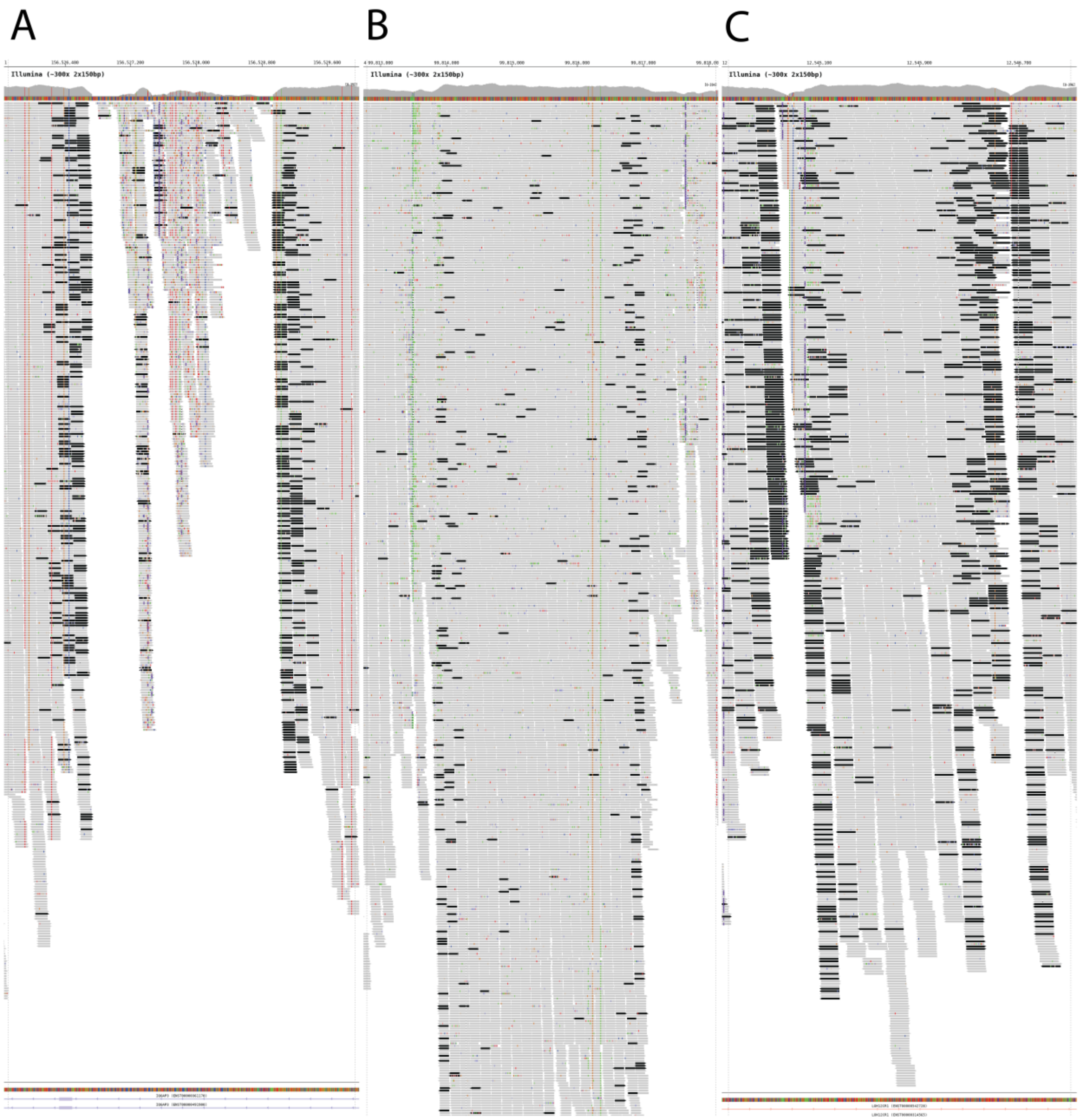


**Supplemental Figure 2. Bamsnap images for multiple region types. A)** The deletion SV from Figure 1 plotted with bamsnap. B) The duplication SV from Figure 2 plotted with bamsnap (cropped to fit). **C)** The inversion SV from Figure 2plotted with bamsnap (cropped to fit).

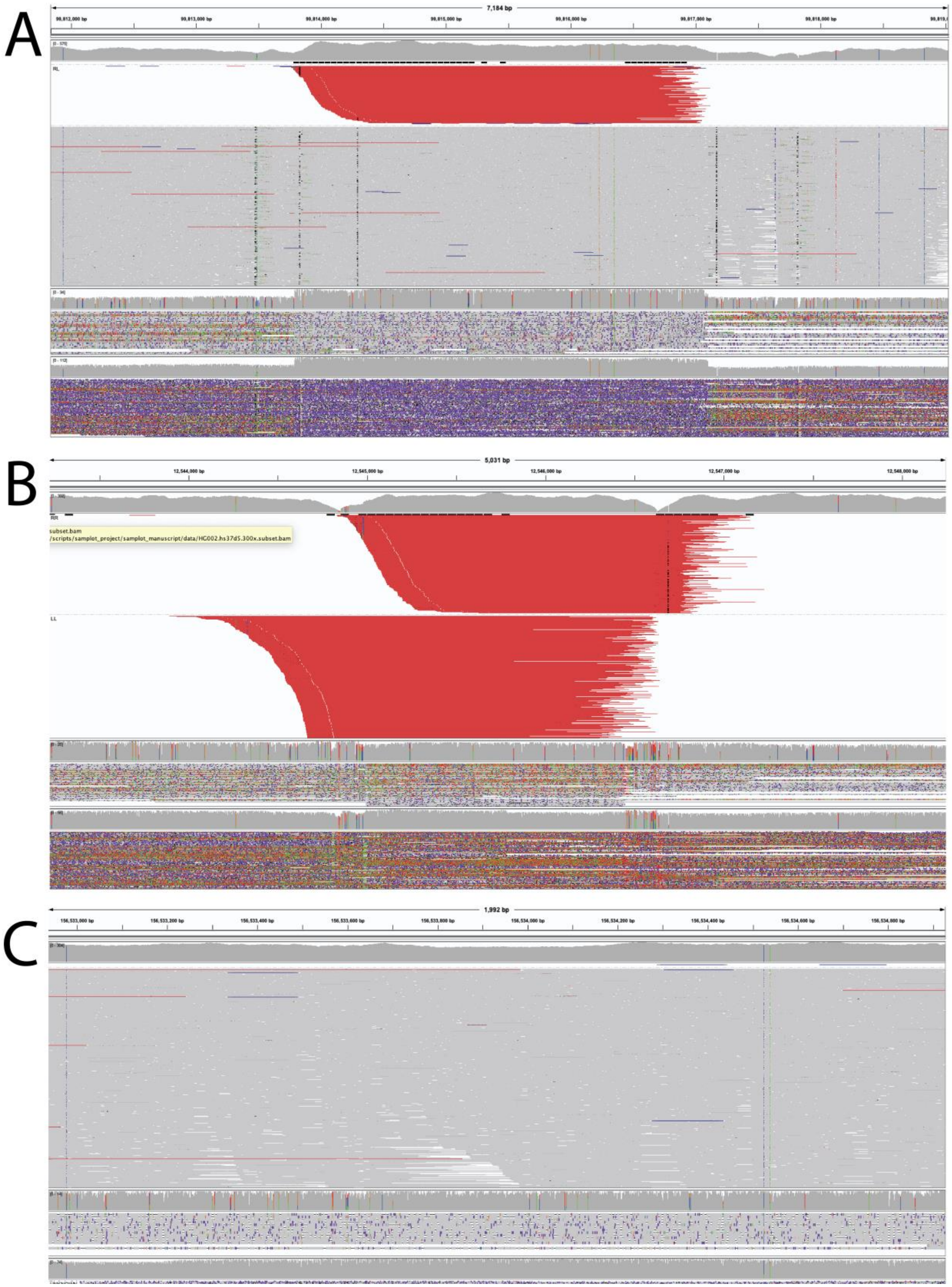


**Supplemental Figure 4. IGV screenshots for multiple region types. A)** The duplication SV from Figure 2 screenshot from IGV. **B)** The inversion SV from Figure 2 screenshot from IGV. **C)** A region with no SV screenshot from IGV.

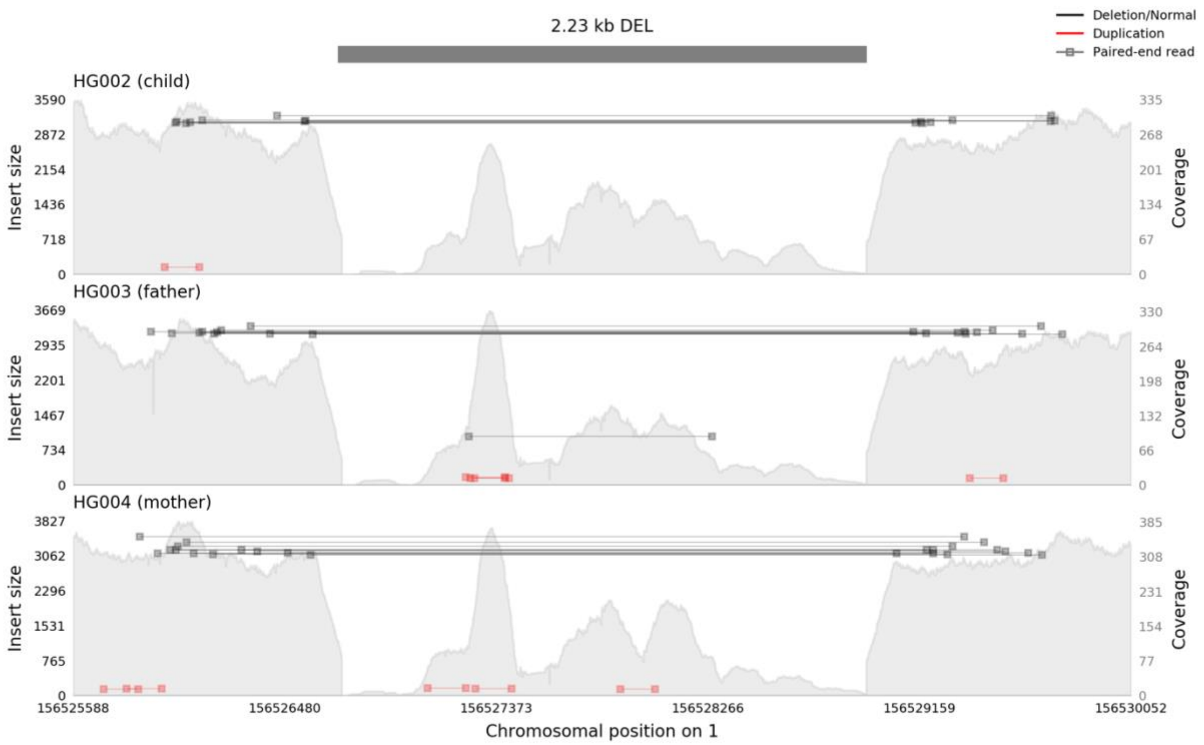


**Supplemental Figure 5. A Samplot image showing a deletion variant in a trio of samples.** Evidence for the variant, in the form of loss of coverage and discordant paired-end reads, appears in the child (top) and both parents.

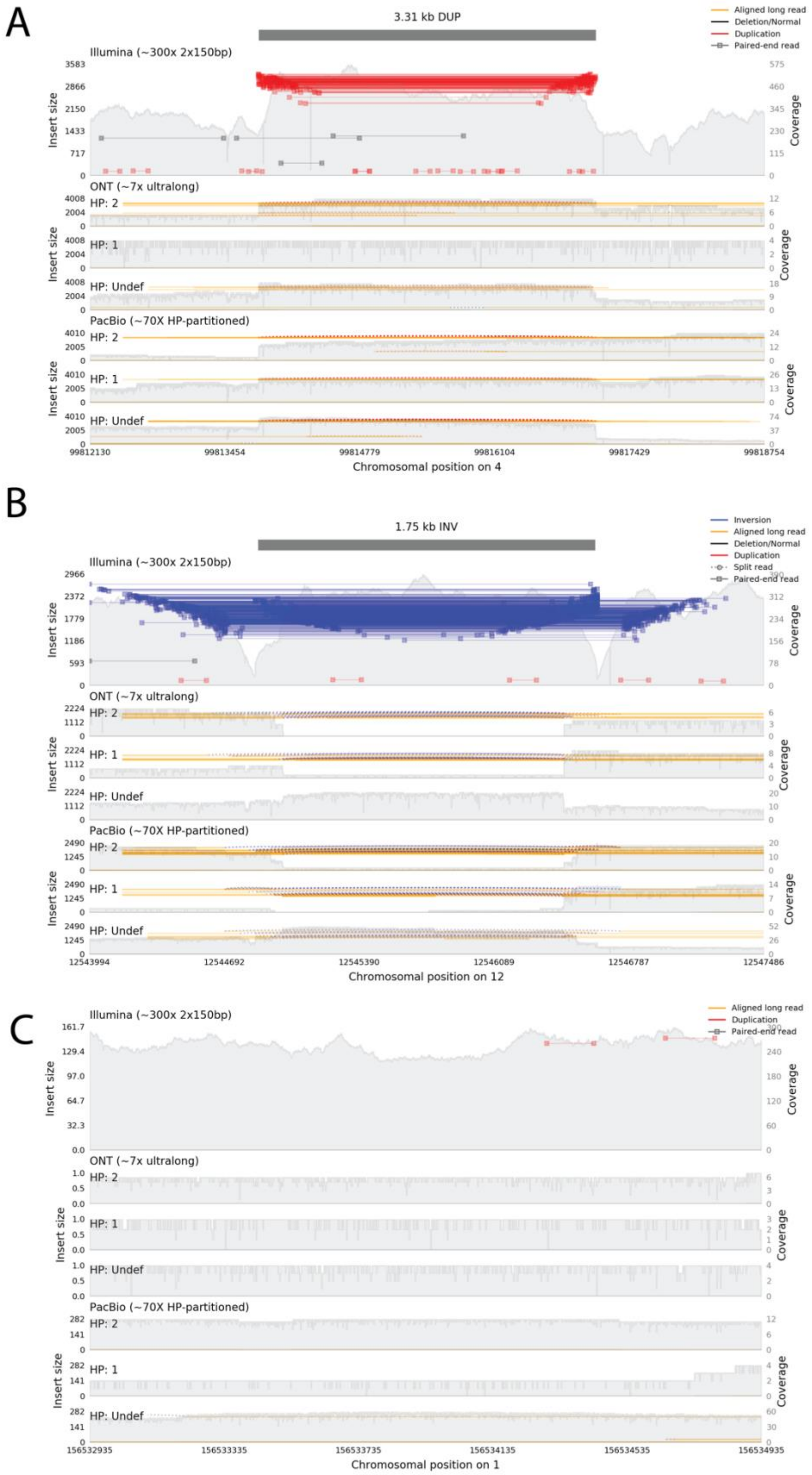


**Supplemental Figure 6. Samplot images of multiple region types with multiple sequencing technologies. A)** The duplication SV from Figure 2 including Illumina, ONT, and PacBio sequence data. **B)** The inversion SV from Figure 2 including Illumina, ONT, and PacBio sequence data. **C)** A region with no SV including Illumina, ONT, and PacBio sequence data.

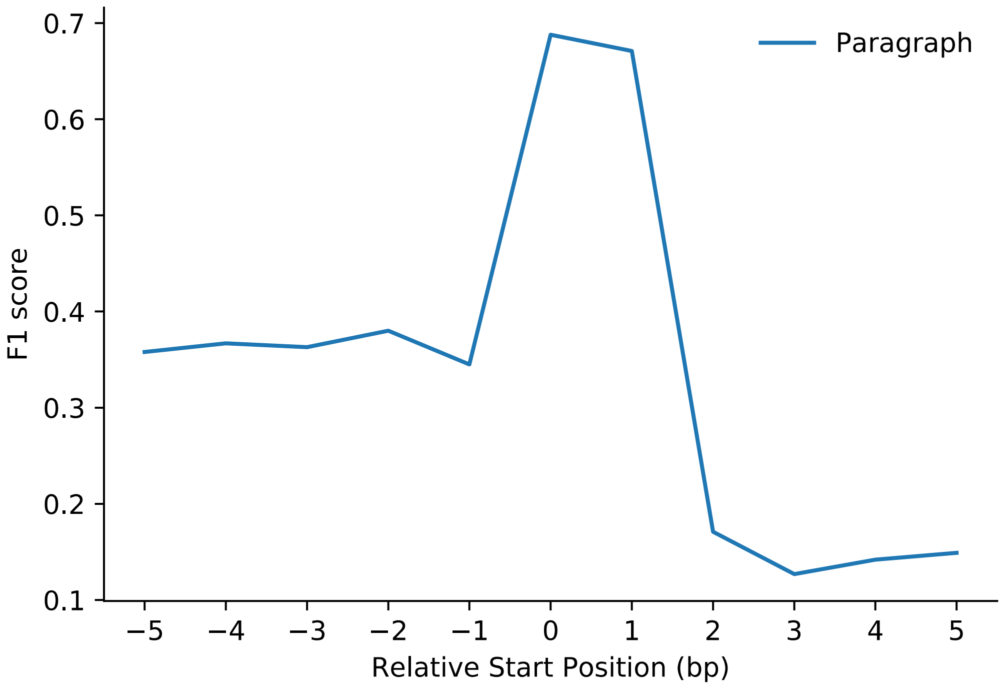


**Supplemental Figure 7. Effect of breakpoint precision on Paragraph genotyping.** Paragraph's genotyping performance, similar to other graph-based methods, is highly sensitive to breakpoint precision. Small deviations in the start position of true positive heterozygous SVs can cause Paragraph to switch genotypes from heterozygous to homozygous reference. This level of sensitivity is problematic because most of the SVs produced by short-read SV callers are not single-base resolution calls. In our experiments, MANTA's average confidence interval size for SV start positions was 8, and LUMPY's was 18

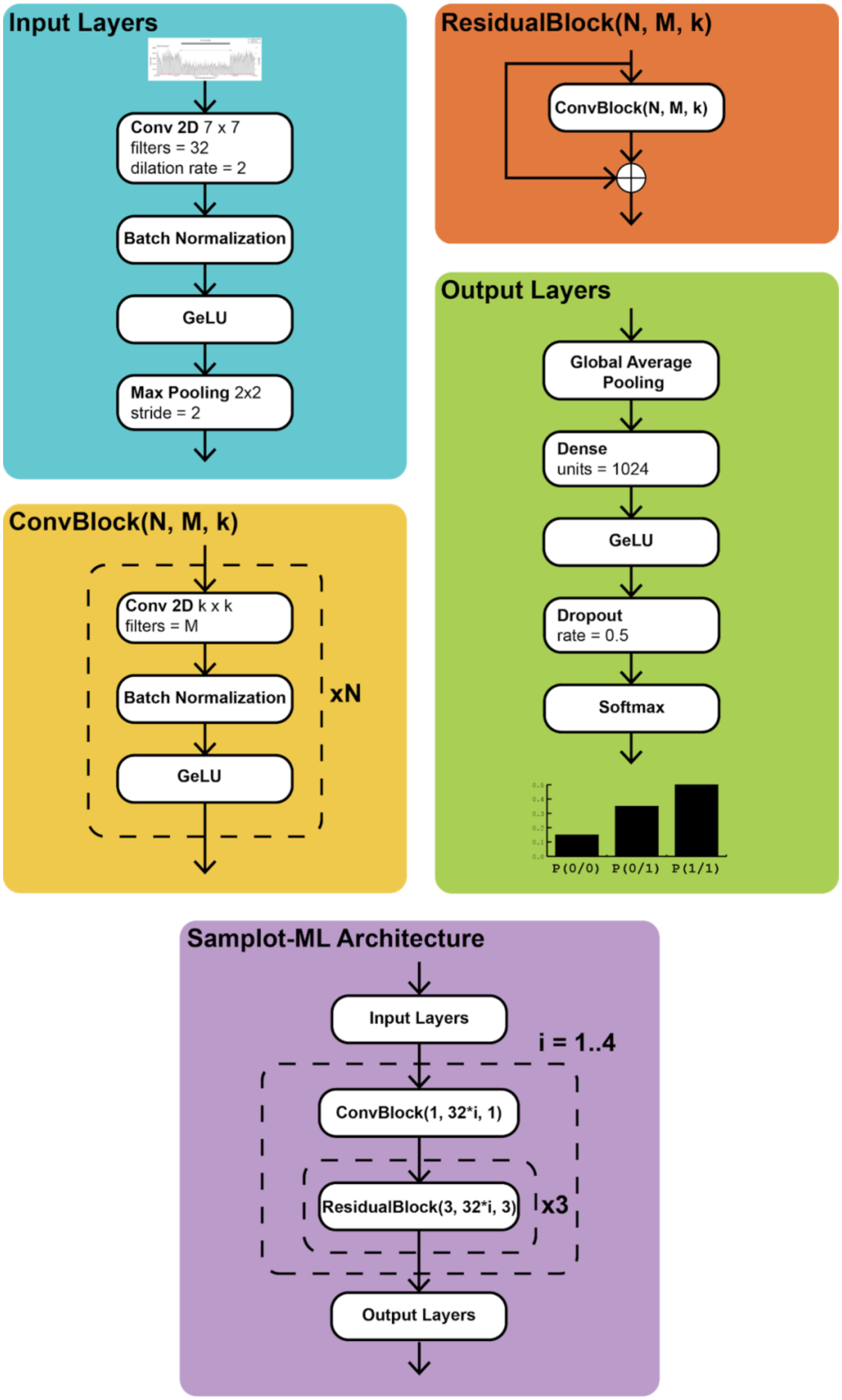


**Supplemental Figure 8. Samplot-ML model architecture**. GeLU refers to the Gaussian error linear unit[34]

#### Running Samplot-ML

From the Samplot-ML root directory:

**1. Generate Images from VCF of deletion SVs**

```
bcftools query -f '%CHROM\t%POS\t%INFO\END\n' $vcf_path |
gargs -p $n_processes \
      "bash data_processing/gen_img.sh \\
      --chrom {0} --start {1} --end {2} \\
      --sample $sample --genotype DEL \\
      --min-mqual 10 \\
      --fasta $fasta_reference \\
      --bam-file $bam \\
      --out-dir $out
```

gargs is an open source alternative to xargs and can be found at https://github.com/brentp/gargs

**2. Crop Images**

```
bash data_processing/crop.sh \
      -p $n_processes \
      -d $path_to_images \
      -o $out
```

**3. Filter input VCF/modify predicted genotypes with Samplot-ML**

```
find $path_to_cropped_images -name '*.png' > image-list.txt
bash evaluation/create_test_vcfs.sh \
      --model-path saved_models/samplot-ml.h5 \
      --data-list image-list.txt \
      --vcf $vcf_path \
      --num-processes $n_processes \
      --batch-size $batch_size \
      --out-dir $out
```

**$n_processes** is the number of cpu processes used to load images to the model. **$batch_size** is the number of images to feed to the model at once. The output directory will contain the filtered vcf and a bed file with regions and prediction scores with format:

```
Chrom    start    end    Pref    Phet    Palt
```

Where Pref, Phet, and Palt are the prediction scores for homozygous reference, heterozygous, and homozygous alternate genotypes.
